# Phase separation promotes Atg8 lipidation for autophagy progression

**DOI:** 10.1101/2024.08.29.610189

**Authors:** Yuko Fujioka, Takuma Tsuji, Tetsuya Kotani, Hiroyuki Kumeta, Chika Kakuta, Toyoshi Fujimoto, Hitoshi Nakatogawa, Nobuo N. Noda

## Abstract

Upon starvation, the autophagy-initiating Atg1 complex undergoes phase separation to organize the pre-autophagosomal structure (PAS) in yeast, from which autophagosome formation is considered to proceed. However, the physiological roles of the PAS as a liquid droplet remain unclear. Here we show that core Atg proteins are recruited into early PAS droplets that are formed by phase separation of the Atg1 complex with different efficiencies in vitro. The Atg12–Atg5–Atg16 E3 ligase complex for Atg8 lipidation is the most efficiently condensed in the droplets via specific Atg12–Atg17 interaction, which is also important for the PAS targeting of the E3 complex in vivo. In vitro reconstitution experiments reveal that E3-enriched early PAS droplets promote Atg8 lipidation and incorporate Atg8-coated vesicles to the interior, thereby protecting them from Atg4-mediated delipidation. These data suggest that the PAS utilizes its liquid-like property to function as an efficient production site for lipidated Atg8 and pool membrane seeds to drive autophagosome formation.

## INTRODUCTION

Phase separation is a general mechanism that reversibly condenses biomolecules such as proteins and nucleic acids in the cytoplasm and the nucleoplasm.^1,2^ The resultant condensates are often called liquid droplets and play crucial roles in various cellular phenomena using the liquid-like property, the typical role of which is to promote or inhibit biochemical reactions by condensing or excluding specific enzymes.^3^ Although phase separation was initially studied as a mechanism of condensing proteins and RNA, recent studies revealed that phase separation is also important for membrane-related phenomena such as clustering synaptic vesicles (SVs) at synapses,^4^ maintaining the integrity of the Golgi ribbon,^5^ and generating autophagosomes in the cytoplasm.^6^ However, the molecular mechanisms of how phase separation drives membrane-related functions remains largely unknown.

Autophagosome is a double membrane-bound organelle that is de novo generated upon induction of autophagy, a lysosomal degradation system important for cellular homeostasis.^7^ In budding yeast, autophagy-related (Atg) proteins are known to be recruited at the pre-autophagosomal structure (PAS), which functions as a site of autophagosome formation.^8^ We recently revealed that the PAS behaves as a liquid droplet.^9^ Upon nutrient starvation, dephosphorylation of Atg13 triggers the formation of the autophagy-initiating Atg1 complex consisting of Atg1, Atg13, Atg17, Atg29, and Atg31, which undergoes phase separation to organize the early PAS droplets by multivalent interactions between Atg13 and Atg17. Early PAS droplets then transform into mature PAS droplets by recruiting downstream Atg proteins that include the phosphatidylinositol (PI)-3 kinase complex consisting of Vps34, Vps15, Vps30/Atg6, Atg14, and Atg38, a lipid transfer protein Atg2, PI 3-phosphate (PI3P) binding proteins Atg18 and Atg21, and the Atg8-phosphatidylethanolamine (PE) conjugation (Atg8 lipidation) system consisting of a ubiquitin-like protein Atg8, the E1 enzyme Atg7, the E2 enzyme Atg3, the E3 enzyme complex consisting of the Atg12–Atg5 protein conjugate and Atg16, the E2 enzyme Atg10 for Atg12–Atg5 conjugation, and the deconjugase Atg4 that delipidates Atg8.^7,10^ Moreover, the PAS droplets recruit a lipid scramblase Atg9-containing vesicles (termed Atg9 vesicles) as an initial membrane seed for generating autophagosomal precursor membranes.^11–13^ However, little is known about the specific roles that PAS droplets play in initial steps of autophagosome formation using their liquid-like properties.

Here, we introduced downstream Atg proteins into early PAS droplets in vitro and demonstrated that PAS droplets function as a site of Atg8–PE production by condensing the E3 complex while excluding the deconjugase. Moreover, we found that PAS droplets incorporate membrane vesicles in an Atg8-lipidation dependent manner, which could function as a mechanism of gathering membrane seeds for initial autophagosomal membrane generation. These data suggest that the PAS droplets drive autophagosome formation by performing a number of direct functions using the liquid-like property.

## RESULTS

### The Atg12–Atg5–Atg16 complex is most efficiently condensed in early PAS droplets in vitro

We previously showed that the Atg1 complex undergoes phase separation to form liquid-like droplets in vitro (termed early PAS droplets hereafter), which function as a scaffold of the PAS assembly in yeast cells.^9^ The mature PAS consists of not only the Atg1 complex, but also other various Atg proteins, which are important for progression of autophagy. We purified core Atg proteins and complexes as follows: 1) the Atg1 complex consisting of Atg1 (kinase-dead D211A mutant), Atg13, Atg17, Atg29, and Atg31; 2) Atg8 and Atg12 conjugation systems including Atg8, Atg4 (processing and deconjugating enzyme for Atg8), Atg7 (E1 for Atg8 and Atg12), Atg3 (E2 for Atg8), the Atg12–Atg5 conjugate complexed with Atg16 (the Atg12–Atg5–Atg16 complex; E3 for Atg8), and Atg10 (E2 for Atg12); 3) the autophagy-specific PI-3 kinase (PI3K) complex consisting of Vps34, Vps15, Atg6/Vps30, Atg14, and Atg38; 4) lipid transfer protein Atg2; and 5) phosphoinositide binding proteins Atg18 and Atg21 (Figure S1). These proteins were fluorescently labeled and individually mixed with pre-formed early PAS droplets and observed by fluorescence microscopy (Figure 1). Intriguingly, efficiency of condensation into early PAS droplets varied widely from protein to protein: the Atg12–Atg5–Atg16 complex showed the highest condensation efficiency, followed by Atg2, Atg21, Atg8, and Atg18. The condensation of Atg3, Atg4, Atg7, Atg10, and PI3K was not significant compared with SNAP or GFP. These data indicate that there is a preference for the protein that early PAS droplets accommodate.

**Figure 1.**
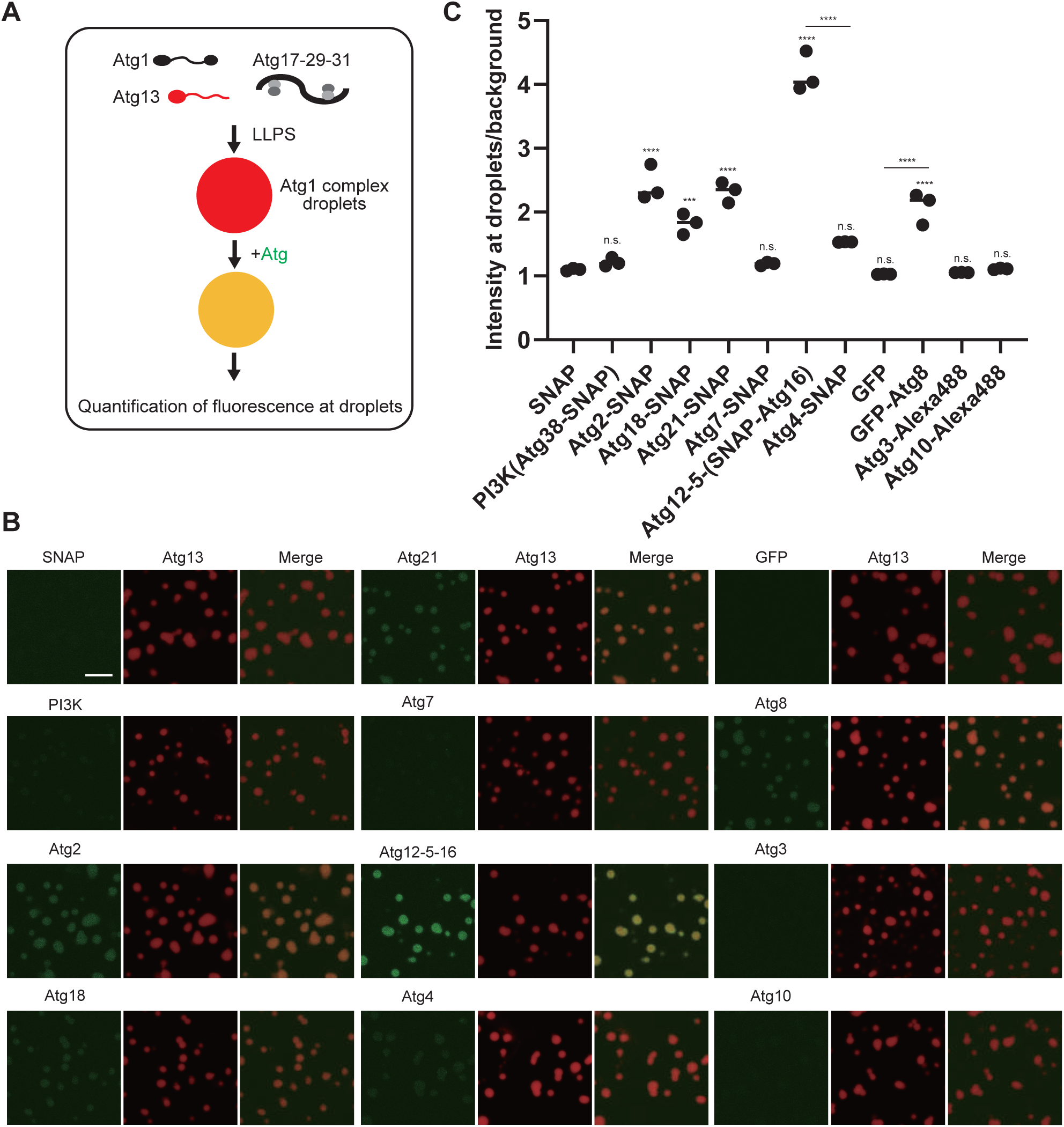
Atg12–Atg5–Atg16 complex is efficiently condensed in early PAS droplets. (A) Schematic drawing of the in vitro experiments. (B) Fluorescence images of early PAS droplets supplemented with each Atg protein. Scale bar = 10 μm. (C) Quantification of (B). \*\*\**P* = 4.4 × 10^−4^ (SNAP vs. Atg18–SNAP), \*\*\*\**P* = 1.4 × 10^−8^ (SNAP vs. Atg2–SNAP), 7.5 × 10^−8^ (SNAP vs. Atg21–SNAP), 1.8 × 10^−14^ (SNAP vs. Atg12–5–16–SNAP), 3.7 × 10^−6^ (SNAP vs. GFP–Atg8), 1.1 × 10^−6^ (GFP vs. GFP–Atg8), n.s., not significant (SNAP vs. PI3K–SNAP, Atg7–SNAP, Atg4–SNAP, GFP, Atg3, and Atg10), using a two-sided Tukey’s multiple comparisons test. See also Figure S1.

### A portion of Atg12 IDR mediates specific interaction with Atg17

We then studied the mechanism of how the Atg12–Atg5–Atg16 complex was efficiently condensed in the droplets. We previously showed that the Atg12–Atg5 conjugate interacts with Atg17 in vivo, for which the N-terminal intrinsically disordered region (IDR) of Atg12 is important.^14^ Moreover, comprehensive protein complex prediction in yeast suggested that Atg12 IDR binds to Atg17.^15^ We used AlphaFold2^16^ to predict the Atg12–Atg17–Atg29–Atg31 complex structure and confirmed that Atg12 IDR, especially residues 42–62, bind to Atg17 without interfering with Atg29–Atg31 binding or homo-dimerization of Atg17 (Figure 2A). In order to predict the detailed interactions, residues 38–63 of Atg12 were used for prediction of Atg17-bound structure (Figure 2B). Residues 42–62 of Atg12 assume an α-helical conformation and bind to the hydrophobic patch of Atg17 consisting of Ile147, Ile154, Tyr303, Ile306, Phe307, Ile310, Leu313, and Phe317 using the side-chains of Val43, Leu47, Phe50, Arg53, Leu54, and Leu57. In addition, hydrophilic interactions were observed between Ser60 and Asp61 of Atg12 and Lys164 and Tyr303 of Atg17 and between Ser51 and Ser55 of Atg12 and Gln157 of Atg17. Hereafter, we call the residues 42–62 of Atg12 as Atg17-binding helix (17BH).

**Figure 2.**
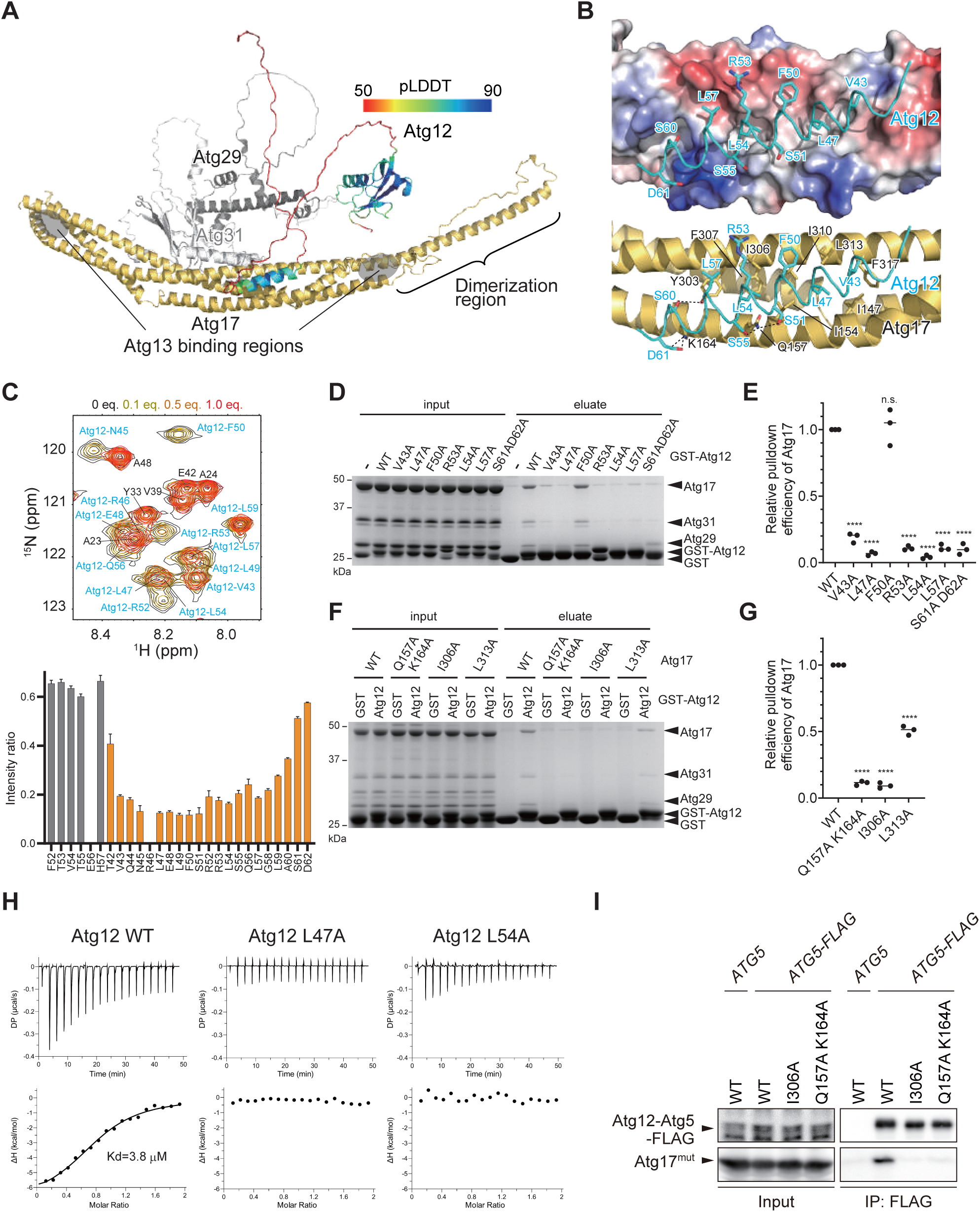
Structural basis of the Atg12–Atg17 interaction. (A) AlphaFold2 structure of Atg12 bound to the Atg17–Atg29–Atg31 complex. Atg17, Atg29, and Atg31 are colored in yellow, dark gray, and light gray, respectively, whereas Atg12 is colored based on the predicted local distance difference test (pLDDT) value. (B) Close-up view of the interactions between Atg12 and Atg17. Top: Atg17 is shown with the surface model with coloring based on the electrostatic potentials (blue and red indicate positive and negative potentials, respectively). Bottom: Atg12 and Atg17 are shown with the ribbon model, in which the side-chains of the residues involved in Atg12–Atg17 interaction are shown with the stick model. Ppossible hydrogen bonds are shown with a broken line. (C) Atg17–Atg31C titration experiment of GB1–Atg12^17BH^ by NMR. Top: a portion was extracted from Figure S2B. GB1-derived signals were labeled in black, and Atg12-derived signals were labeled in blue. Bottom: the signal intensity ratio before and after the addition of 0.5 molar equivalents of Atg17-Atg31C was extracted from Figure S2C, and GB1 is shown in gray and Atg12 in orange. The signal-to-noise ratio of the signals was used to calculate the error bars. (D) In vitro pulldown assay between GST–Atg12 mutants and Atg17–Atg29– Atg31. (E) Quantification of (D). \*\*\*\**P* = 6.4 × 10^−11^ (WT vs. L43A), 9.6 × 10^−12^ (WT vs. L47A), 1.7 × 10^−11^ (WT vs. R53A), 6.5 × 10^−12^ (WT vs. L54A), 1.9 × 10^−11^ (WT vs. L57A), 1.6 × 10^−11^ (WT vs. S61A D62A), n.s., not significant (WT vs. F50A), using a two-sided Tukey’s multiple comparisons test. (F) In vitro pulldown assay between GST–Atg12 and Atg17–Atg29–Atg31 mutants. (G) Quantification of (F). \*\*\*\**P* = 1.1 × 10^−10^ (WT vs. Q157A K164A), 7.8 × 10^−11^ (WT vs. I306A), 1.5 × 10^−8^ (WT vs. L313A), using a two-sided Tukey’s multiple comparisons test. (H) ITC results were obtained by titration of the GB1–Atg12^17BH^ into a solution of the Atg17–Atg31C complex. (I) Coimmunoprecipitation of Atg5–FLAG and Atg17 mutants from yeast cells treated with rapamycin for 2 h. See also Figure S2.

We then performed NMR analyses on Atg12^17BH^ fused to a GB1 tag, which showed that Atg12^17BH^ has a partial α-helical conformation and that the NMR signals of Atg12^17BH^ were much more severely attenuated than those of the GB1 moiety upon Atg17 titration (Figures 2C and S2). These data suggest that Atg12^17BH^ is actually involved in the interaction with Atg17, presumably in the α-helical conformation. In vitro pulldown assay using GST-fused Atg12^17BH^ and Atg17 confirmed their direct interaction, and this interaction was severely impaired by Ala-substitution at most of the residues of Atg12 and Atg17 that were predicted to be involved in the Atg12–Atg17 interaction (Figures 2D-2G). We then analyzed the interaction using isothermal titration calorimetry and showed that the binding affinity between Atg12^17BH^ and Atg17 was 3.8 μM, which was severely impaired by Ala substitution at Leu47 or Leu54 of Atg12 (Figure 2H). We further analyzed the interaction of the Atg12–Atg5 conjugate with Atg17 by coimmunoprecipitation experiments and confirmed that the Atg17 residues important for Atg12^17BH^ binding in vitro are also important for the interaction in vivo (Figure 2I). These results indicate that the predicted Atg12–Atg17 interaction is sufficiently reliable and that Atg12^17BH^ forms specific direct interaction with Atg17 both in vitro and in vivo.

### Specific Atg12–Atg17 interaction is necessary and sufficient for being condensed in early PAS droplets

We next studied whether the specific interaction between Atg12^17BH^ and Atg17 is responsible for the condensation of the Atg12–Atg5–Atg16 complex in early PAS droplets. We first confirmed that the condensation of the E3 complex in the droplets was significantly impaired by deletion of Atg12 IDR and more severely impaired by deletion of whole Atg12 from the E3 complex (Figures 3A and 3B), indicating that Atg12 IDR is necessary for this process. Although GFP alone was not condensed, fusion of Atg12^17BH^ to GFP dramatically promoted its condensation in the droplets, indicating that Atg12^17BH^ functions as a targeting sequence to early PAS droplets (Figure 3C). Furthermore, Ala substation at Val43, Leu47, Leu54, or Leu57 in Atg12^17BH^ severely impaired the condensation of the fusion protein in the droplets (Figures 3C and S3A). Taken together, these data suggest that the specific Atg12–Atg17 interaction as predicted by AlphaFold2 is necessary and sufficient for the condensation of the Atg12–Atg5–Atg16 complex in early PAS droplets.

**Figure 3.**
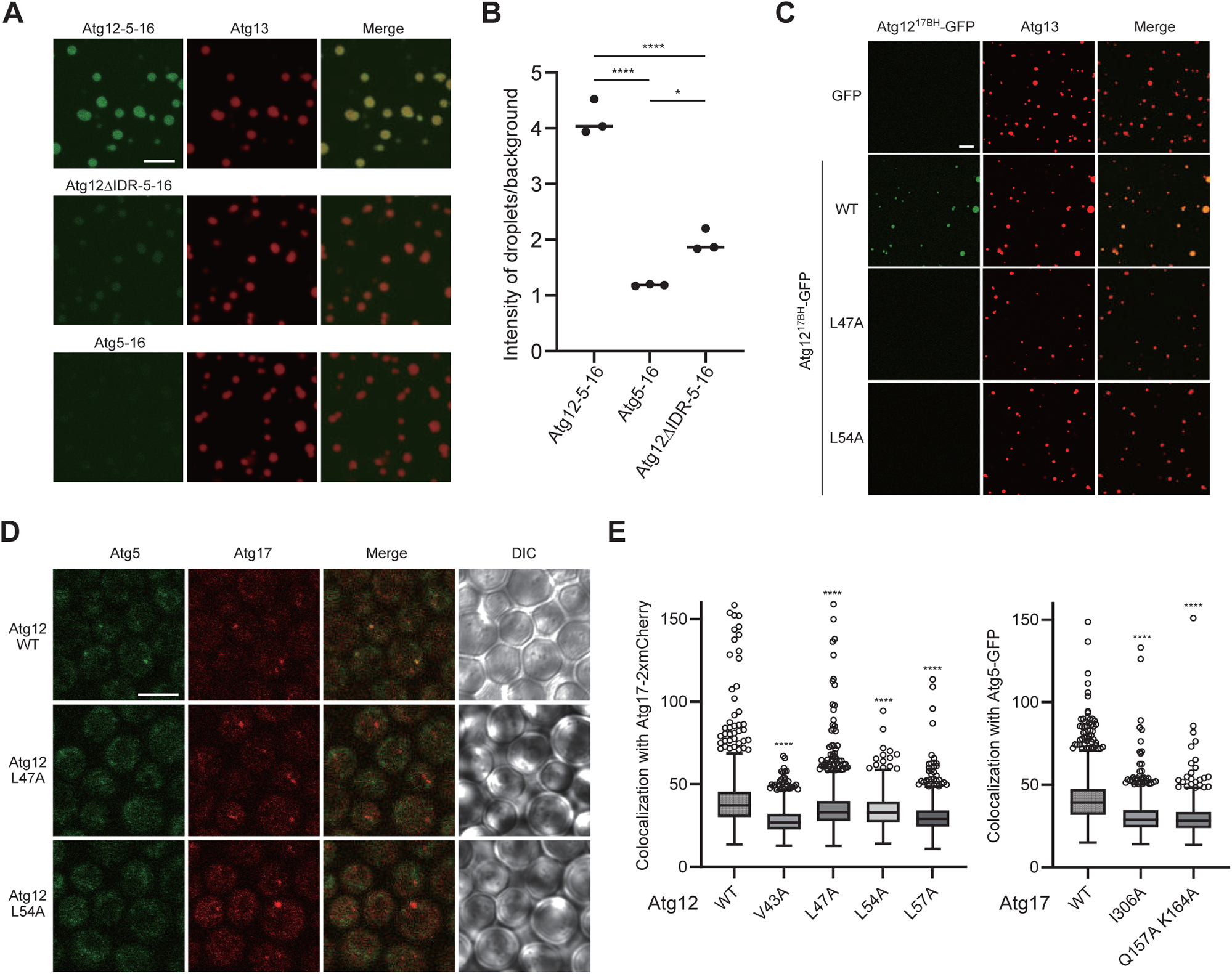
Atg12–Atg17 interaction is responsible for the in vitro and in vivo condensation of the E3 complex in the droplet. (A) Fluorescence images of the early PAS droplets supplemented with each Atg protein. Atg12–5–16 images are the same as those used in Figure 1B. Scale bar = 10 μm. (B) Quantification of fluorescence intensity of SNAP–Atg16 in the droplets in (A). \**P* = 1.04 × 10^−2^, \*\*\*\**P* = 6.3 × 10^−6^ (Atg12-5-16 vs. Atg5-16), 4.0 × 10^−5^ (Atg12–5–16 vs. Atg12ΔIDR–5–16), using a two-sided Tukey’s multiple comparisons test. (C) Fluorescence images of the early PAS droplets supplemented with GB1– Atg12^17BH^. Scale bar = 10 μm. (D) Fluorescence images of GFP–Atg5 and mCherry–Atg17 in yeast cells. Scale bar = 5 μm. (E) Quantification of the colocalization of GFP–Atg5 with mCherry–Atg17 at the PAS puncta in (D). Number of PAS puncta: n = 1,548 (WT), 1,990 (V43A), 1,815 (L47A), 1,710 (L54A), and 1,829 (L57A). \*\*\*\**P* < 1.0 × 10^−15^, using a two-sided Dunn’s multiple comparisons test. See also Figure S3.

### Specific Atg12–Atg17 interaction mediates Atg21-independent PAS targeting of the Atg12–Atg5–Atg16 complex in vivo

We previously reported that the Atg12–Atg5–Atg16 complex targets to the PAS via two distinct mechanisms, one is via the Atg16–Atg21–PI3P interaction and another via the Atg12–Atg5 conjugate–Atg17 interaction. We speculated that the specific Atg12–Atg17 interaction we characterized here is responsible for the latter mechanism. We observed the PAS localization of the Atg12–Atg5–Atg16 complex by monitoring GFP–Atg5 in *atg12*Δ *atg17*Δ *atg21*Δ cells co-expressing Atg5–GFP, Atg12, and Atg17-2xmCherry. Colocalization of GFP–Atg5 with Atg17–2xmCherry was observed in cells expressing wild-type Atg12 and Atg17, which was significantly reduced in cells expressing wild-type Atg17 and Atg12 mutants (V43A, L47A, L54A, or L57A) or expressing wild-type Atg12 and Atg17 mutants (Q157A K164A or I306A) (Figures 3D, 3E, S3B, and S3C). These data, together with the binding data, suggest that the Atg12–Atg5–Atg16 complex targets to the PAS via specific Atg12–Atg17 interaction in the absence of Atg21.

### The Atg1 complex and Atg21 cooperatively promote lipidation and inhibit delipidation of Atg8 in vitro

The findings that the early PAS droplet specifically condenses the Atg12–Atg5–Atg16 complex, the E3 enzyme for Atg8 lipidation, led us to consider the possibility that it could act as a reaction site for Atg8 lipidation. We then performed in vitro Atg8 lipidation reaction using Atg8, Atg7 (E1), Atg3 (E2), Atg12–Atg5– Atg16 (E3), liposomes mimicking the lipid composition of Atg9 vesicles (phosphatidylcholine (PC):PI:PE:phosphatidylserine (PS):phosphatidic acid (PA)=41:44:7:7:2),^17^ and MgATP in the presence or absence of early PAS droplets, which were confirmed to be formed in this reaction condition (Figure 4A). We also studied the effect of Atg21 on Atg8 lipidation because Atg21 was reported to recruit the Atg12–Atg5–Atg16 complex to PI3P-containing membranes via binding to both Atg16 and PI3P and promote Atg8 lipidation.^18,19^ Consistent with previous reports, Atg21 promoted Atg8 lipidation in vitro although statistically not significant (Figures 4B, 4C, and S4A). Importantly, early PAS droplets promoted Atg8 lipidation more efficiently and statistically significantly, the promoting activity of which was also detected in the presence of Atg21 (Figures 4B and 4C). These data suggest that early PAS droplets promote Atg8 lipidation irrespective of the presence or absence of Atg21.

**Figure 4.**
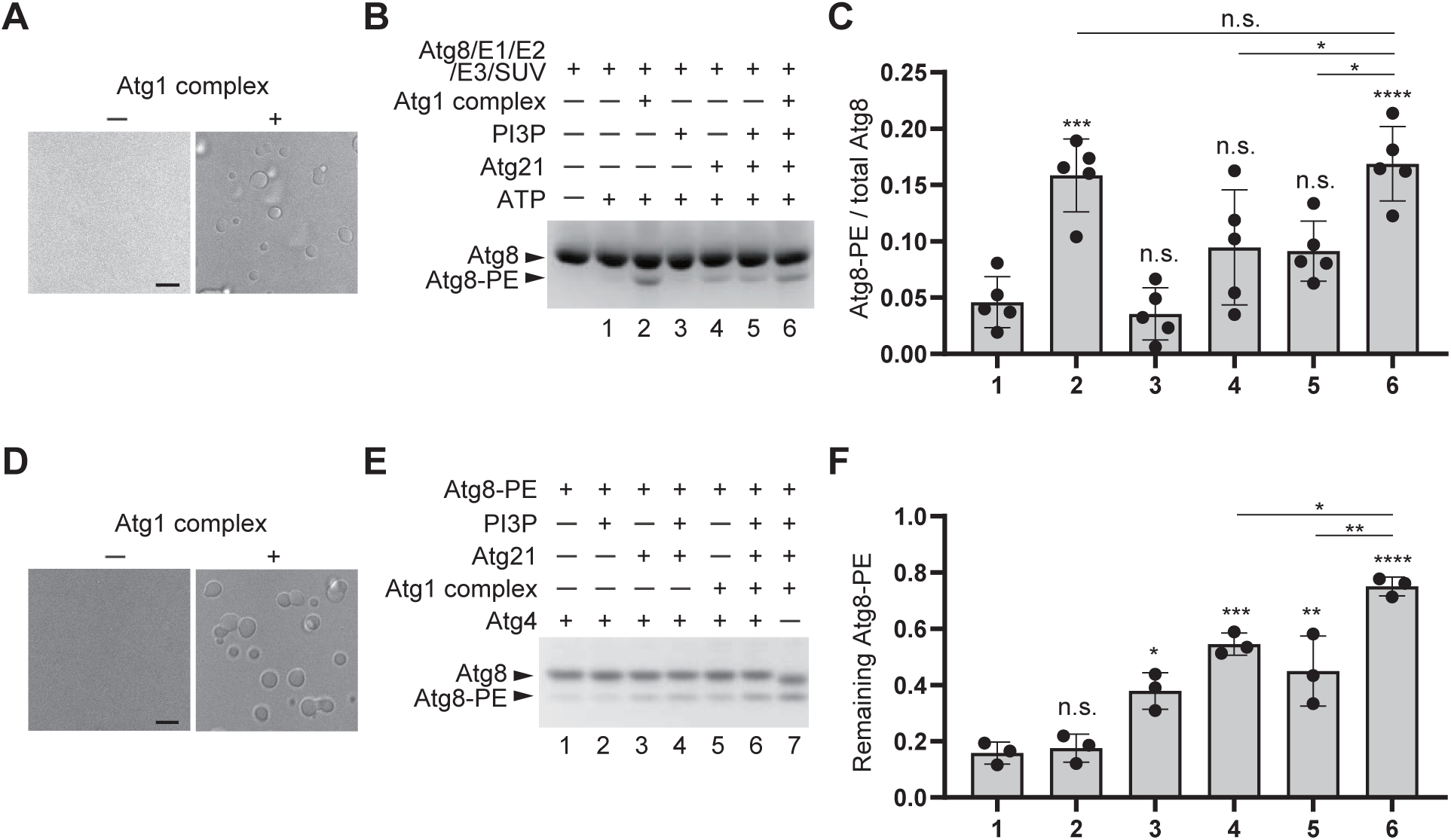
Early PAS droplets promote Atg8 lipidation and inhibit Atg8–PE delipidation in vitro. (A) Observation of the early PAS droplets in the reaction solution using differential interference contrast microscopy. The left and right images correspond to reaction conditions 1 and 2 in (B), respectively. (B) Effect of the early PAS droplets, Atg21, and PI3P on the Atg8 lipidation reaction. The reaction solutions were subjected to urea-SDS-PAGE, followed by Coomassie brilliant blue staining. (C) Quantification of the ratio between Atg8–PE and total Atg8 in (B). (D) Observation of the early PAS droplets in the reaction solution using differential interference contrast microscopy. The left and right images correspond to reaction conditions 1 and 5 in (E), respectively. (E) Effect of the early PAS droplets, Atg21, and PI3P on the Atg8–PE delipidation reaction. The reaction solutions were subjected to urea-SDS-PAGE, followed by Coomassie brilliant blue staining. (F) Quantification of the ratio between Atg8–PE and total Atg8 in (E). Each value was divided by the value in lane 7 (without Atg4). See also Figure S4.

We then studied the effect of early PAS droplets and Atg21 on the efficiency of Atg8 delipidation by Atg4 in vitro. After performing Atg8 lipidation reaction using E1, E2, E3, and MgATP on liposomes, the reaction was stopped by addition of dithiothreitol and further supplemented with Atg4 and either early PAS droplets (which were confirmed to be formed at this reaction condition, Figure 4D), Atg21, or buffer solution (Figure S4B). As shown in Figures 4E and 4F, the band of Atg8–PE disappeared upon incubation with Atg4, indicating that the used Atg4 was active and delipidated Atg8 in this reaction condition. Addition of early PAS droplets or Atg21 significantly inhibited the delipidation reaction by Atg4. Addition of both early PAS droplets and Atg21 further severely inhibited the delipidation reaction. These data suggest that early PAS droplets and Atg21 additively inhibit Atg8 delipidation reaction in vitro.

### Specific Atg12–Atg17 interaction is important for Atg8 lipidation and autophagy in the absence of Atg21

We then studied the roles of the specific Atg12–Atg17 interaction in Atg8 lipidation and autophagy activity in yeast cells. Because the Atg16–Atg21 interaction redundantly functions in the recruitment of the Atg12–Atg5–Atg16 complex to the PAS,^14^ we used *atg12*Δ strains with or without *atg21* deletion. In both *atg12*Δ and *atg12*Δ *atg21*Δ strains, Atg8–PE was not observed at all, which was recovered by expression of wild-type Atg12 (Figures 5A and 5B). Expression of Atg12 mutants that are defective in Atg17-binding could also recover Atg8–PE formation in the case of *atg12*Δ strains, whereas it failed to recover Atg8–PE formation in the case of *atg12*Δ *atg21*Δ strains. These data suggest that the Atg12–Atg17 interaction is crucial for Atg8 lipidation in the absence of Atg21 in vivo. We also measured bulk autophagy activity by monitoring the processing of Pgk1–EGFP, a non-selective autophagy cargo that releases free EGFP upon delivery to the vacuole via autophagy.^20^ In both *atg12*Δ and *atg12*Δ *atg21*Δ strains, free EGFP was not observed at all, which was recovered by expression of wild-type Atg12 (Figures 5C and 5D). Expression of Atg12 mutants that are defective in Atg17-binding could also recover EGFP processing in the case of *atg12*Δ strains, whereas it failed to recover EGFP processing in the case of *atg12*Δ *atg21*Δ strains. These data suggest that the Atg12–Atg17 interaction is crucial for autophagy activity in the absence of Atg21, which is consistent with the levels of Atg8–PE.

**Figure 5.**
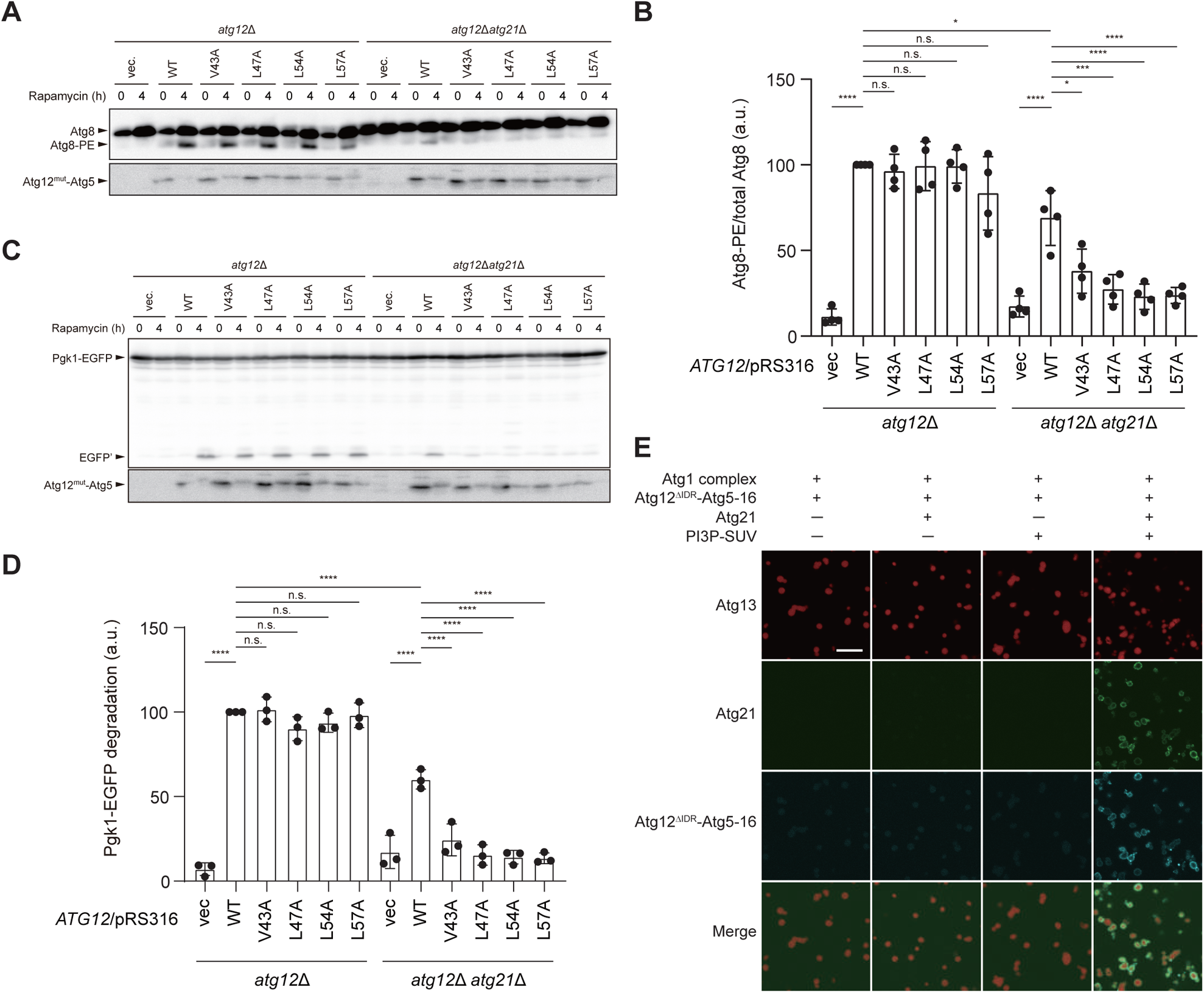
Atg12–Atg17 interaction is important for autophagy in the absence of Atg21. (A) Observation of Atg8–PE accumulation in yeast cells. (B) Quantification of the ratio between Atg8–PE and total Atg8 in (A). \**P* = 1.6 × 10^−2^ (WT in *atg12*Δ vs. WT in *atg12*Δ *atg21*Δ), 1.7 × 10^−2^ (WT vs. V43A in *atg12*Δ *atg21*Δ), \*\*\**P* = 3.5 × 10^−4^ (WT vs. L47A in *atg12*Δ *atg21*Δ), \*\*\*\**P* = 1.5 × 10^−11^ (WT vs. vec in *atg12*Δ), 7.9 × 10^−6^ (WT vs. vec in *atg12*Δ *atg21*Δ), 6.9 × 10^−5^ (WT vs. L54A in *atg12*Δ *atg21*Δ), 9.3 ×10^−5^ (WT vs. L57A in *atg12*Δ *atg21*Δ), n.s., not significant, using a two-sided Tukey’s multiple comparisons test. (C) Pgk1–EGFP processing assay. (D) Quantification of Pgk1–EGFP degradation in (C). \*\*\*\**P* < 1.0 × 10^−12^ (WT vs. vec in *atg12*Δ), \*\*\*\**P* = 3.7 × 10^−6^ (WT in *atg12*Δ vs. WT in *atg12*Δ *atg21*Δ), 9.8× 10^−7^ (WT vs. vec in *atg12*Δ *atg21*Δ), 2.2 × 10^−5^ (WT vs. V43A in *atg12*Δ *atg21*Δ), 4.6 × 10^−7^ (WT vs. L47A in *atg12*Δ *atg21*Δ), 2.9 × 10^−7^ (WT vs. L54A in *atg12*Δ *atg21*Δ), 2.2 × 10^−7^ (WT vs. L57A in *atg12*Δ *atg21*Δ), n.s., not significant, using a two-sided Tukey’s multiple comparisons test. (E) In vitro Atg21 and PI3P-dependent targeting of the Atg12ΔIDR–Atg5–Atg16 complex in the early PAS droplets observed by fluorescence microscopy. Scale bar = 10 μm. See also Figure S5.

In vivo analyses showed that Atg8–PE is sufficiently formed without the specific Atg12–Atg17 interaction in the presence of Atg21. We then studied the Atg21-mediated recruitment of the E3 complex to early PAS droplets in vitro (Figures 5E and S5). Consistent with Figure 3A, lack of Atg12^IDR^ severely, and lack of Atg12 more severely impaired the condensation of the E3 complex in early PAS droplets. Atg21 alone was not efficiently localized at early PAS droplets and failed to promote the recruitment of the E3 complex lacking Atg12 or its IDR. On the other hand, PI3P-containing liposomes interacted with the surface of early PAS droplets and promoted the targeting of Atg21 to the droplet surface, and importantly, the combination of Atg21 and PI3P-containing liposomes recruited the E3 complex lacking Atg12 or its IDR efficiently to the droplet surface. These data suggest that Atg21 and PI3P-containing liposomes together can recruit the E3 complex to early PAS droplets without the Atg12–Atg17 interaction.

### Atg8 lipidation promotes incorporation of lipid vesicles into early PAS droplets in vitro

We finally observed the behavior of liposomes with lipid compositions mimicking Atg9 vesicles during the Atg8 lipidation reaction in the presence of early PAS droplets using freeze fracture replica electron microscopy, in which deep etching was performed so that the protein droplets embedded in ice were exposed (Figure S6A; the procedure is summarized in Figure 6A). Then PS, a component of the liposomes, as well as Atg8 was immune-stained. As shown in Figures 6B and 6C, PS and Atg8 signals were accumulated inside the droplets only when incubated with MgATP. We also detected PS signals inside the droplets when Atg8 lipidation reaction was performed in the presence of Atg21 and PI3P-SUVs but in the absence of the Atg12–Atg17 interaction (Figure S6B). These data clearly demonstrate that liposomes are condensed within early PAS droplets by Atg8 lipidation via either the Atg12–Atg17 interaction or the Atg16–Atg21 interaction pathway (Figure 7).

**Figure 6.**
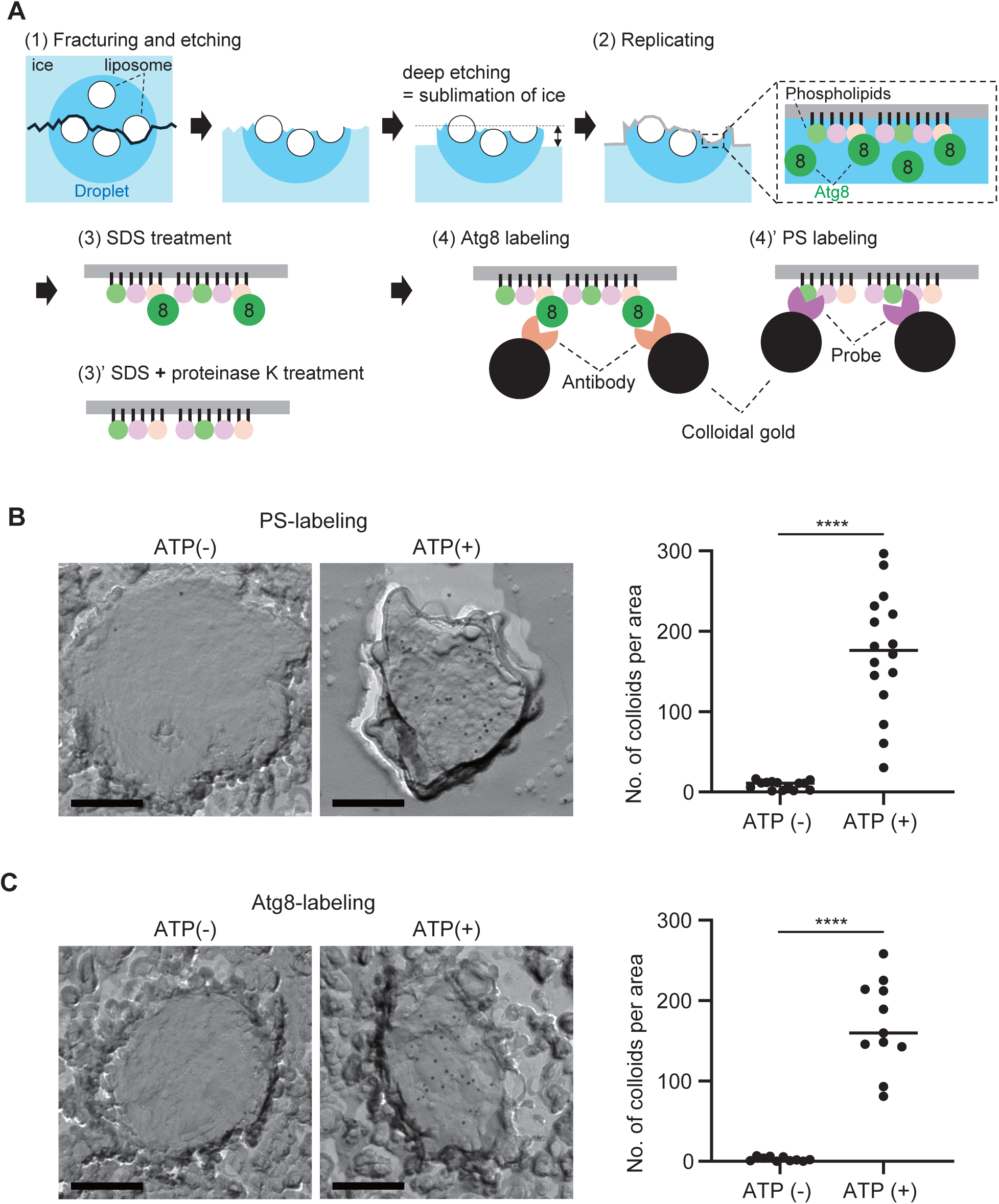
Early PAS droplets incorporate Atg8-coated vesicles via two distinct pathways. (A) Schematic drawing of the procedures for freeze-fracture replica electron microscopy. (B) Atg8 lipidation-dependent detection of the phosphatidylserine signals in the Atg1 complex droplet observed by freeze-fracture replica electron microscopy. The right graph shows the number of colloids per area of each droplet. Number of droplets analyzed: n = 14 for ATP (-) and 16 for ATP (+). \*\*\*\**P* = 1.4 × 10^−8^, using a two-sided Mann-Whitney test. Scale bar = 200 nm. (C) Atg8 lipidation-dependent detection of Atg8 signals in the Atg1 complex droplet observed by freeze-fracture replica electron microscopy. The right graph shows the number of colloids per area of each droplet. Number of droplets analyzed: n = 10 for ATP (-) and 11 for ATP (+). \*\*\*\**P* = 5.7 × 10^−6^, using a two-sided Mann-Whitney test. Scale bar = 200 nm. See also Figure S6.

**Figure 7.**
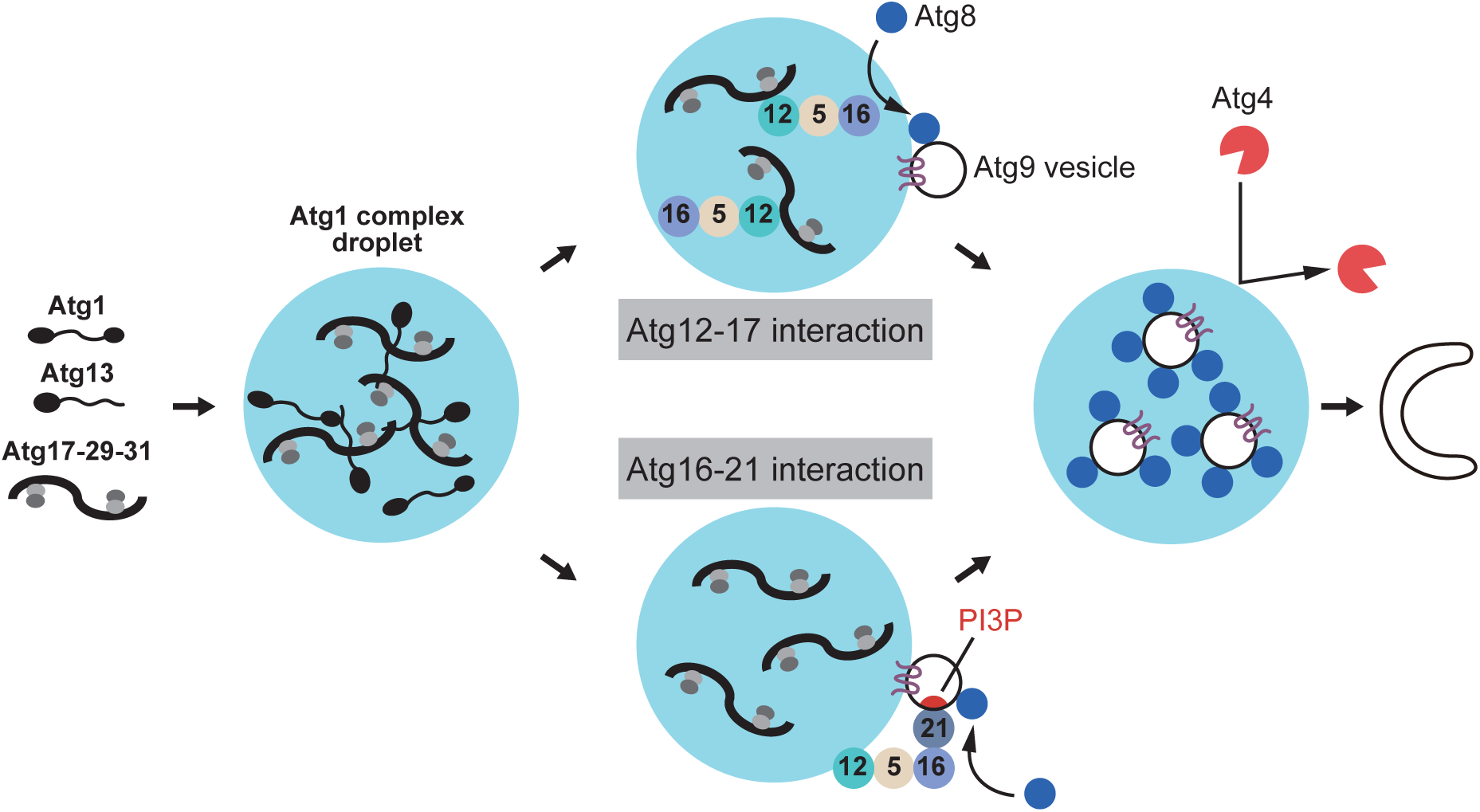
Proposed model of the processes in the PAS droplet during autophagy initiation.

## Discussion

In budding yeast, starvation induces phase separation of the Atg1 complex into early PAS droplets, which lead to the construction of the mature PAS and initiate autophagosome formation.^9^ However, it remained elusive why the PAS needs to be a liquid-like droplet and what molecular roles the PAS exerts using the liquid-like property. Here, we show that the early PAS droplets condense the Atg12– Atg5–Atg16 E3 complex for Atg8 lipidation via the specific interaction of the Atg12^17BH^ with Atg17 while eliminating Atg4, the deconjugase for Atg8, thereby promoting Atg8 lipidation while inhibiting Atg8 delipidation in vitro. The latter inhibitory function of the droplet can explain the previous in vivo observation that Atg4 is responsible for deconjugating Atg8–PE produced at the organelles other than the PAS with unknown mechanisms.^21–23^ This dual function of the PAS as a liquid droplet will enable local production of sufficient amounts of Atg8–PE for autophagy progression. A role of phase separation in promotion of ubiquitination has been reported for HECT-type E3 Itch.^24^ It will be interesting to see if the dual function of phase separation observed for the Atg8 conjugation system is also used in ubiquitin and other ubiquitin-like conjugation systems.

As the earliest event of autophagosome formation, it has been proposed that the PAS droplet recruits Atg9 vesicles and utilizes them as membrane seeds for generating initial autophagic membranes.^9,11^ However, this process leaves many open issues. Even the precise localization of Atg9 vesicles at the PAS droplet, whether it is the surface or interior, has not been determined due to the small size of the PAS. Using the reconstitution system, we clearly demonstrate that the early PAS droplets internalize membrane vesicles mimicking the lipid composition and size of Atg9 vesicles in the Atg8 lipidation-dependent manner (Figures 6B and 6C), proposing that Atg9 vesicles would also localize at the interior of the PAS droplet upon autophagy induction. This phenomenon is reminiscent of the incorporation of SVs into phase separated synapsin condensates at the presynapse.^4,25^ Synapsin condensates selectively incorporate SVs but not other organelles such as mitochondria and endoplasmic reticulum despite their localization in close proximity. This selectivity could be attributed to the small size of SVs (∼45 nm), their characteristic high content of negatively charged lipids (∼20 mol%), or the membrane proteins such as VAMP2 and synaptophysin embedded in SVs, although the actual mechanism is still unclear.^4^ The mechanism by which the early PAS droplets incorporate Atg9 vesicles, which are similar to SVs in size (30-60 nm) and in their high content of negatively charged lipids (∼53 mol%), presumably with selectivity, is also not yet fully understood. However, the dependency of liposomes on Atg8 lipidation for incorporation into early PAS droplets observed in vitro gives us a substantial hint for the mechanism. One possible idea is that some specific interactions of Atg8 conjugated on the vesicles with the Atg1 complex components could be a mechanism for vesicle internalization. Actually, Atg8 alone showed modest condensation efficiency into early PAS droplets (Figure 1). Alternatively, the attachment of positively charged Atg8 (calculated PI value is 8.9) to the negatively charged membrane surface could promote wetting of the vesicles with the early PAS droplets that are negatively charged (calculated PI value of the Atg1 complex is 5.6) by promoting electrostatic interactions.

One role of the condensation of Atg9 vesicles within the PAS droplets is to protect Atg8–PE produced on the vesicles from Atg4 by spatial separation, which is supported by in vitro deconjugation assay (Figures 4D-4F). Then, are there any other roles for Atg9 vesicle condensation in the PAS droplets? It is known that enforced contact between lipid vesicles promotes their fusion in vitro.^26^ It could be speculated that Atg9 vesicles trapped within narrow PAS droplets are forced to interact closely with each other, which promote their homotopic fusion to generate initial autophagic membranes in concert with PAS-localizing factors such as Atg8–PE, which is known to perturb membranes and promote their tethering and hemi/full-fusion in vitro.^27–30^ The reconstitution study presented here provides the basis for unlocking these secrets and leading to a complete understanding of the mechanisms of autophagosome formation.

## METHODS

### Construction of expression plasmids

Expression plasmids encoding Atg1^D211A^, Atg17–Atg29–Atg31, and Atg17– Atg31C (residues 174–196) were constructed as previously described.^31^ An expression plasmid encoding N-terminal His-tagged and C-terminal SNAP-Twin-Strep-tagged Atg13 was constructed as previously described.^9^ The expression plasmids encoding the processed form of Atg8^K26P^, Atg7, Atg3, and Atg10 were constructed as previously described.^32–35^ Expression plasmids encoding the Atg12–Atg5 conjugate, Atg12ΔIDR (residues 100–186)–Atg5 conjugate, and Atg5 were constructed as previously described.^36,37^ The expression plasmids encoding the Atg7–SNAP, Atg4 (residues 28–494)–SNAP, SNAP–Atg16 were constructed as previously described.^28,38^

To construct the expression plasmids encoding N-terminal GST-tagged Atg12^17BH^, N-terminal GST-tagged SNAP, N-terminal GST-tagged mGFP, N-terminal GST-tagged and C-terminal SNAP-tagged Atg38, N-terminal GST-tagged and C-terminal mGFP-tagged Atg12^17BH^, and N-terminal GST and mGFP-tagged Atg8^K26P^, the genes were amplified by polymerase chain reaction (PCR) and cloned into pGEX6p-1 vectors. To construct an expression plasmid encoding N-terminal GB1-His-tagged Atg12^17BH^, the gene was amplified by PCR and cloned into a pGBHPS vector.^39^ To construct an expression plasmid encoding the C-terminal SNAP-tagged Atg21 with an HRV3C protease recognition site followed by a His tag and a SUMO tag, the genes were amplified by PCR and cloned into a pE-SUMOpro Amp vector. To construct yeast expression plasmids encoding Atg2 and Atg18, respectively, followed by Twin-Strep, SNAP, and His tags, the genes were amplified by PCR and cloned into a pRS426-based multi-copy plasmid under the control of the GAL1 promoter. The cloning procedure was performed using the NEBuilder HiFi DNA Assembly Cloning Kit (New England Biolabs). The mutations that resulted in the indicated amino acid substitutions were introduced via PCR-mediated site-directed mutagenesis. To confirm the identity of the constructs, they were all sequenced.

### Protein expression and purification

Atg1^D211A^, Atg17-Atg29-Atg31, Atg17-Atg31C, and their mutants with the amino acid mutations indicated in the main text were purified as previously described.^31^ N-terminal His-tagged and C-terminal SNAP–Twin-Strep-tagged Atg13 was purified as previously described.^9^ The processed forms of Atg8^K26P^, Atg7, Atg3, and Atg10 were purified as previously described.^32–35^ Atg7–SNAP, Atg4 (residues 28–494)–SNAP, SNAP–Atg16, and Atg12–Atg5–SNAP–Atg16 were purified as previously described.^28,38^ Atg12ΔIDR–Atg5–SNAP–Atg16 and Atg5–SNAP– Atg16 were purified in a similar manner as that of the full-length or non-tagged proteins, respectively.^38,40^

To express the recombinant proteins in *Escherichia coli*, the BL21(DE3) strain was used as the expression host. GST–Atg38–SNAP was purified using a COSMOGEL® GST-Accept column (Nacalai). Subsequently, the GST was excised with human rhinovirus 3C (HRV3C) protease, and the eluate was desalted using a Bio-Rad Bio-Gel P-6 Desalting Cartridge (Bio-Rad). Subsequently, Atg38–SNAP was applied to a GST-Accept column to remove the excised GST. The expression of the PI3K complex I (composed of Vps34, Vps30, Vps15, and Atg14) was performed as previously described.^41^ During the incubation of the lysate with immunoglobulin G (IgG) beads, Atg38–SNAP was added to form the PI3K complex I pentamer. Subsequent purification steps were conducted in a similar manner as those of a previously described method.^12^

The GST–SNAP, GST–mGFP, GST–mGFP–Atg8K26P, GST–Atg12 (residues 42–62), and GST–Atg12 (residues 42–62)–mGFP were initially purified using a GST-Accept column. Subsequently, the samples were desalted using a Bio-Gel P-6 desalting cartridge. Phosphate-buffered saline (PBS) was used as the buffer. For the purification of GST–SNAP, GST–mGFP, GST–mGFP–Atg8^K26P^, and GST–Atg12^17BH^–mGFP, GST was excised using HRV3C protease. Subsequently, the samples were subjected to another GST-Accept column purification step, to remove the excised GST. Finally, the protein was purified on a HiLoad 26/60 Superdex 200 PG column (Cytiva), with elution performed using 20 mM HEPES [pH 7.0] and 150 mM NaCl.

Atg21–SNAP–HisSUMO was purified via affinity chromatography with a Ni-NTA column (Qiagen). After affinity chromatography, the HisSUMO tag at the C-terminus was excised with HRV3C protease. Subsequently, Atg21–SNAP was purified on a HiLoad 26/60 Superdex 200 PG column eluted with 20 mM HEPES [pH 7.0] and 250 mM NaCl.

Atg2 and Atg18 expression were performed using BJ3505 yeast cells, which lack endogenous Atg2 and Atg18 genes.^42^ The yeast cells harboring the Atg2–Twin-Strep–SNAP–His expression plasmid were cultured in a SD + CA + AT medium (0.17% yeast nitrogen base without amino acids and ammonium sulfate, 0.5% ammonium sulfate, 0.5% casamino acids, and 2% glucose with 0.002% adenine sulfate and 0.002% tryptophan at 30°C. The expression of the protein was induced by the addition of galactose at a final concentration of 4%. The subsequent purification was performed as previously described,^42^ except for the following amendments: The sample that had been eluted from the HisPur Ni-NTA resin (Thermo Fisher Scientific) and subsequently dialyzed was applied to the Strep-Tactin® XT 4Flow® high-volume resin (IBA LifeSciences). The resin was then washed with buffer ST, which consisted of 100 mM Tris-HCl [pH 8.0], 500 mM NaCl, and 10% glycerol. Following elution from the resin with buffer ST containing 50 mM biotin, the eluate was subjected to dialysis against a buffer consisting of 20 mM HEPES [pH 7.0] and 250 mM NaCl. Similarly, Atg18–Twin-Strep–SNAP–His was expressed and prepared. The yeast cells were suspended in a solution of 100 mM Tris-HCl [pH 8.0] and 500 mM NaCl. After the addition of phenylmethylsulfonyl fluoride (PMSF) to the lysate at a final concentration of 1 mM, the yeast cells were disrupted with zirconia beads, centrifuged, and avidin was added to the resulting supernatant at a final concentration of 50 μg/mL. The lysate was then applied to a Strep-Tactin XT high-capacity resin column. Following the resin washing step, Atg18–Twin-Strep–SNAP–His was eluted using a buffer comprising 100 mM Tris-HCl [pH 8.0], 500 mM NaCl, and 50 mM biotin. The protein was then subjected to a HiLoad 26/60 Superdex 200 PG column and eluted with 20 mM HEPES [pH 7.0] and 150 mM NaCl.

The SNAP-tags of Atg2, Atg4, Atg7, Atg16, Atg18, Atg21, and Atg38 were labeled with SNAP-Surface Alexa Fluor 488 (New England BioLabs), the SNAP-tag of Atg13 was labeled with SNAP-Surface 549 (New England BioLabs), and the SNAP-tag of Atg16 for Figure 5E was labeled with SNAP-Surface 647 (New England BioLabs), in accordance with the manufacturer’s instructions. Atg13-SNAP used in Figure 1B was supplemented with the SNAP-Surface Block (New England BioLabs) after labeling.

Atg3 and Atg10 were chemically labeled with Alexa Fluor 488 C5-Maleimide (Thermo Fisher) in a 2:1 molar ratio. The mixtures were incubated on ice for more than 1 h, after which they were applied to a PD SpinTrap™ G-25 (Cytiva) to exclude any residual dye.

### Immunoprecipitation

The *Saccharomyces cerevisiae* strains used in this study are listed in Table S1.^43^ Cells carrying the pRS316-derived plasmids expressing Atg17 and its mutants were cultured in SD + CA + AT to a mid-log phase, converted to spheroplasts, and incubated at 30°C in 0.5 × SD + CA + AT containing 1 M sorbitol and 0.2 µg/mL rapamycin for 2 h. The spheroplasts were solubilized in IP buffer (50 mM Tris-HCl [pH 8.0], 150 mM NaCl, 10% glycerol, 5 mM EDTA, 5 mM EGTA, and 50 mM NaF) containing 2 mM PMSF, 2×cOmplete Protease Inhibitor Cocktail (Roche), and 1% n-dodecyl-β-maltoside (DDM). The samples were clarified by centrifugation at 2,000 ×*g* for 5 min. The supernatants were further centrifuged at 15,000 ×*g* for 15 min. The resulting supernatants (input) were incubated with NHS FG-beads (Tamagawa Seiki) conjugated with anti-FLAG M2 antibody (F1804; Sigma-Aldrich) at 4°C for 2 h while rotating. The beads were washed three times with IP buffer containing 0.1% DDM. The bound proteins were eluted with SDS sample buffer (100 mM Tris-HCl [pH 7.5], 2% SDS, 10% glycerol, and a trace amount of bromophenol blue) at 65°C for 10 min. After removing the beads, DTT was added to the eluates to a final concentration of 20 mM. The samples were analyzed by immunoblotting using antibodies against Atg5 and Atg17.^44,45^

### Pgk1–EGFP cleavage assay

Yeast cells harboring the pRS316-derived plasmids expressing Atg12 and its mutants were cultured in SD + CA + AT to the mid-log phase and treated with rapamycin. Samples for immunoblotting analysis were prepared as previously described.^46^ For immunoblotting, antibodies against GFP (11814460001; Roche) and Atg5 were used.

### Analysis of Atg8–PE formation

Yeast cells harboring the pRS316-derived plasmids expressing Atg12 and its mutants were cultured in SD + CA + AT to the mid-log phase, treated with 1 mM PMSF for 10 min, and then treated with rapamycin. Samples for immunoblotting analysis were prepared as previously described.^46^ Lipidated Atg8 (Atg8–PE) was separated from the unmodified form by urea-sodium dodecyl sulfate-polyacrylamide gel electrophoresis (SDS-PAGE)^47^ and detected using antibodies against Atg8 (anti-Atg8-2).^21^

### Imaging of yeast cells

Yeast cells were cultured until the cell density reached the optical density (OD)600 range of 4.0–6.0. Rapamycin (Sigma-Aldrich) was then added to the cell culture medium to a final concentration of 0.2 μg/mL, and the cells were cultured for an additional 3 h to induce autophagy. The cells were imaged on a glass bottom dish (Mattek) with an FV4000RS confocal laser scanning microscope (EVIDENT) equipped with a UPLXAPO60XO, NA 1.42 Oil objective (EVIDENT) or an FV3000RS confocal laser scanning microscope (EVIDENT) equipped with a UPLSAPO60XO, NA 1.42 Oil objective (EVIDENT).

### Liposome preparation

Soy PI (#840044C), dioleoylphosphatidylethanolamine (PE) (#850725C), 1,2-dioleoyl-sn-glycero-3-phosphocholine (PC) (#850375C), 18:1 dioleoylphosphatidylserine (PS) (#840035C), 1,2-dioleoyl-sn-glycero-3-phosphate (PA) (#840875C), and 18:1 PI3P (#850150P) were purchased from Avanti Polar Lipids. Phospholipids were mixed in a glass vial at the appropriate ratios in chloroform and dried with rotation under a nitrogen gas stream. Samples were dried further in a desiccator under vacuum for 16 h. The resulting lipid film was suspended in a buffer (20 mM Tris-HCl [pH 7.5] and 150 mM NaCl) at a final concentration of 3 mM phospholipid by vigorous mixing. Samples were then subjected to sonication for 5 min to obtain small unilamellar liposomes.

### Reconstitution and microscopy of protein-rich droplets

Liquid droplets of the Atg1 complex were formed by the dilution of proteins from a stock solution into the buffer as previously described,^9^ with minor modifications. Briefly, liquid droplets of 4 μM Atg1 complex (Atg1^D211A^, Atg13–SNAP, Atg17– Atg29–Atg31) were formed by the dilution of proteins from a stock solution into the buffer (50 mM HEPES [pH 7.0] and 250 mM NaCl) with subsequent incubation at 25°C for 5 min. Then 0.5 μM of each Atg protein/complex was added to the liquid droplets and incubated at 25°C for 15 min for Figure 1B and 3A; 3 μM of Atg12^17BH^–mGFP was added to the liquid droplets and incubated at 25 °C for 5 min for Figure 3C; 0.3 μM of Atg5–SNAP–Atg16 or Atg12ΔIDR–SNAP– Atg16 with or without Atg21–SNAP and 0.5 mM of PI3P-containing liposomes (PC:PE:PI:PS:PA:PI3P at a ratio of 41.1:6:39.3:7:1.6:5 (mol%)) were added to the liquid droplets and incubated at 25 °C for 15 min for Figure 5E. These samples were mixed in a microtube and imaged on a glass bottom dish (Mattek) coated with 3% bovine serum albumin (BSA) (Wako).

These experiments, except for Figure 5E, were performed using an FV3000RS confocal laser scanning microscope (Olympus) equipped with a UPLSAPO60XO, NA 1.42 Oil objective (Olympus). The experiments for Figure 5E were performed using a STELLARIS 5 confocal laser scanning microscope (Leica) equipped with an HC PL APO CS2 63x, NA 1.40 Oil objective (Leica). Quantitative analysis was performed with FIJI.

### In vitro pulldown assay

The purified proteins were incubated with GST-Accept beads at 4°C for 30 min. After three washes with PBS, the proteins were eluted using 10 mM glutathione in 50 mM Tris-HCl [pH 8.0]. The samples were separated by SDS-PAGE. The protein bands were then detected by One Step CBB (BIO CRAFT).

### Isothermal titration calorimetry analysis

Atg12^17BH^–mGFP and Atg17–Atg31C were prepared in the same buffer, which consisted of 20 mM HEPES [pH 7.0] and 150 mM NaCl. The binding of Atg12^17BH^–mGFP to Atg17–Atg31C was measured by ITC using a MICROCAL PEAQ-ITC calorimeter (Malvern) with stirring at 750 rpm. at 25°C. The titration of Atg12^17BH, WT^–mGFP with Atg17–Atg31C involved 18 injections of 2 μL of the Atg12^17BH, WT^–mGFP solution (220 μM) at intervals of 150 s into a sample cell containing 200 μL of Atg17–Atg31C (22 μM). The titration of the Atg12^17BH, L47A^– mGFP with Atg17–Atg31C involved 18 injections of 2 μL of the Atg12^17BH, L47A^–mGFP solution (224 μM) at intervals of 150 s into a sample cell containing 200 μL of Atg17–Atg31C (21 μM). The titration of the Atg12^17BH, L54A^-mGFP with Atg17–Atg31C involved 18 injections of 2 μL of the Atg12^17BH, L54A^–mGFP solution (232 μM) at intervals of 150 s into a sample cell containing 200 μL of Atg17– Atg31C (23 μM). The raw titration data were analyzed using the MicroCal PEAQ-ITC Analysis Software with a single-site binding model. The error of each parameter shows the fitting error.

### Atg8 lipidation assay

Liquid droplets of the Atg1 complex were formed by the dilution of proteins from a stock solution into the buffer as previously described, with minor modifications. ^9^ Briefly, liquid droplets of 4 μM of the Atg1 complex (Atg1^D211A^, Atg13–SNAP, Atg17–Atg29–Atg31) were formed by the dilution of proteins from a stock solution into the buffer (50 mM HEPES [pH 7.0] and 250 mM NaCl) with subsequent incubation at 30 °C for 10 min. During incubation, purified 2 μM of Atg7, 2 μM of Atg3, 0.2 μM of Atg12–Atg5–Atg16, 20 μM of the C-terminal glycine-exposed form of Atg8^K26P^, 2 μM of Atg21, and 0.75 mM of liposomes were combined in a reaction buffer (20 mM HEPES [pH 7.0], 150 mM NaCl, and 2 mM DTT). After the droplet formation reaction was complete, it was mixed with the lipidation reaction solution in equal volumes, to which 1 mM ATP and 1 mM MgCl^2^ were added and incubated at 30°C for 5 min. The reaction was stopped by boiling in the SDS-PAGE sample buffer. The samples were subjected to urea-SDS-PAGE and stained with One Step CBB (BIO CRAFT).

### Assay for Atg8–PE delipidation

Purified 1 μM of Atg7, 1 μM of Atg3, 0.1 μM of Atg12–Atg5–Atg16, 10 μM of the C-terminal glycine-exposed form of Atg8^K26P^, and 0.38 mM of liposomes were combined in a reaction buffer (20 mM HEPES [pH 7.0], 150 mM NaCl, 1 mM DTT, 1 mM ATP, and 1 mM MgCl^2^) with subsequent incubation at 30°C for 10 min. After the lipidation reaction was complete, the reaction was stopped by adding 100 mM of DTT. Liquid droplets of the Atg1 complex were formed as described above. The droplets, 1 μM of Atg4, and 1 μM of Atg21 were mixed in equal volumes with the lipidation reaction solution and incubated at 30°C for 15 min. The reaction was stopped by boiling in the SDS-PAGE sample buffer. The samples were subjected to urea-SDS-PAGE and stained with One Step CBB.

### Quick-freezing and freeze-fracture replica labeling

Liquid droplets of 4 μM of the Atg1 complex (Atg1^D211A^, Atg13–SNAP, Atg17– Atg29–Atg31) were formed by the dilution of proteins from a stock solution into the buffer (50 mM HEPES [pH 7.0] and 250 mM NaCl) with subsequent incubation at 25 °C for 10 min. The solution was then mixed with Atg8 lipidation reaction solution (purified 2 μM of Atg7, 2 μM of Atg3, 2 μM of Atg12-Atg5-Atg16, 20 μM of the C-terminal glycine-exposed form of Atg8K26P, and 0. 75 mM liposomes in a reaction buffer consisting of 20 mM HEPES [pH 7.0], 150 mM NaCl, and 2 mM DTT with or without 1 mM ATP and 1 mM MgCl^2^) in equal volumes and further incubated at 25°C for 20 min. The specimens were quick-frozen using an HPM 010 high-pressure freezing machine according to the manufacturer’s instructions (Leica Microsystems, Wetzlar, Germany). The frozen specimens were transferred to the cold stage of ACE900 (Leica Microsystems, Wetzlar, Germany), fractured at −102°C under a vacuum of ∼1 × 10^−6^ mbar, and maintained at the same temperature for 3 min to induce sublimation of water from the fractured surface. Replicas were made by electron-beam evaporation in three steps: (1) carbon (6 nm thick) at a 90° angle to the specimen surface; (2) platinum-carbon (2 nm thick) at a 45° angle; and (3) carbon (10 nm thick) at a 90° angle. For PS labeling, thawed replicas were treated with 2.5% SDS in 0.1 M Tris-HCl [pH 8.0] containing 50 μg/mL proteinase K (Nacalai, Kyoto, Japan) at 60°C overnight. Replicas were then incubated at 4°C overnight with GST-tagged 2× evectin-2 PH domain (150 ng/mL) in 0.1 M phosphate buffer [pH 6.0] containing 1% BSA and 1% cold-water fish gelatin, followed by rabbit anti-GST antibody (5 μg/mL, Bethyl Laboratories, Texas, USA) and colloidal gold (10 nm)-conjugated protein A (PAG10; University of Utrecht, Utrecht, The Netherlands), for 40 min each at 37°C in 1% BSA in PBS.^48^ For Atg8 labeling, replicas were treated with 2.5% SDS in 0.1 M Tris-HCl [pH 8.0] at 60°C overnight and incubated with rabbit anti-Atg8-IN13 antibody^27^ (kindly provided by Dr. Yoshinori Ohsumi of the Tokyo Institute of Technology), followed by PAG10. The labeled replicas were observed using a JEOL JEM-1400plus transmission electron microscope (Tokyo, Japan).

### NMR experiments

All NMR experiments were performed using a Bruker 800 MHz NMR AVANCE NEO with a CPTCI probe or a Bruker 600 MHz NMR AVANCE III HD with a TBI probe. The measurement temperature was 25°C. The NMR sample used for the main chain assignment was a ^13^C/^15^N-labeled 1.1 mM GB1–Atg12 sample, adjusted to pH 7.0 with 20 mM HEPES buffer containing 150 mM NaCl. The following standard spectral sets were used for main chain assignment; HNCO, HNCA, CBCA (CO)NH, HBHA (CO)NH, ^15^N-NOESY (mix 150 msec). As a result, we were able to assign all main chain amide signals observed on [^1^H-^15^N] HSQC. All signals derived from GB1 and Atg12 were assigned (Figure S2A). The secondary structure propensities (SSP)^49^ were calculated from the obtained chemical shift value information (Figure S2D). ^15^N-labeled GB1–Atg12 adjusted to 0.1 mM was used for the titration experiments. Atg17–Atg31C adjusted to their respective ratios (0, 0.1, 0.5, and 1.0 molar equivalents) were added, and the [^1^H-^15^N] HSQC was measured (Figure S2B). The signal intensity ratio before and after the addition of Atg17-Atg31C was plotted (Figure S2C).

## ACKNOWLEDGMENTS

We thank Yohei Ohashi and Roger L Williams for providing the plasmids to purify the PI3K proteins; Hayashi Yamamoto for providing the plasmids for expressing the Atg17 mutants; Yuki Ishikawa, Konomi Marumo, Sonoko Kurazono, and Elijah William Caldwell for assistance with the protein preparation; Ayaka Saito for assistance with EM sample preparation; and Yuta Ogasawara for assistance with fluorescence imaging quantification. This work was supported in part by JSPS KAKENHI Grant Number: JP21H05731, JP23H02429, JP23H04923, JP23K27122 to Y.F.; JP22K06818, JP22H04654 to T.T.; JP22K06123 to T.K.; JP22H00446 to T.F.; JP19H05708 to H.N.; JP23K20044 to H.N. and N.N.N.; JP19H05707, JP24H00060 to N.N.N.; PRIME, Japan Agency for Medical Research and Development Grant number JP20gm6410009 to Y.F.; CREST, Japan Science and Technology Agency Grant number JPMJCR20E3 to N.N.N.; and grants from the Takeda Science Foundation to Y.F. and N.N.N.

## AUTHOR CONTRIBUTIONS

Y.F. and N.N.N. conceived the project. Y.F. performed *in vitro* experiments. T.T. and T.F. performed replica electron microscopy. H.K. performed NMR analysis. Y.F., T.K., C.K. and H.N. performed yeast experiments. All authors analyzed the data. Y.F. and N.N.N. wrote the manuscript with input from all other authors. Y.F. and N.N.N. supervised the work.

## Lead contact

Further inquiries and requests for materials should be directed to the lead contact, Nobuo N. Noda.

## Materials availability

All requests for resources and reagents including plasmids and cell lines are available from the lead contact subject to a Materials Transfer Agreement.

