## Supplemental Figures for "Phase separation promotes Atg8 lipidation for autophagy progression"

#### Figure S1

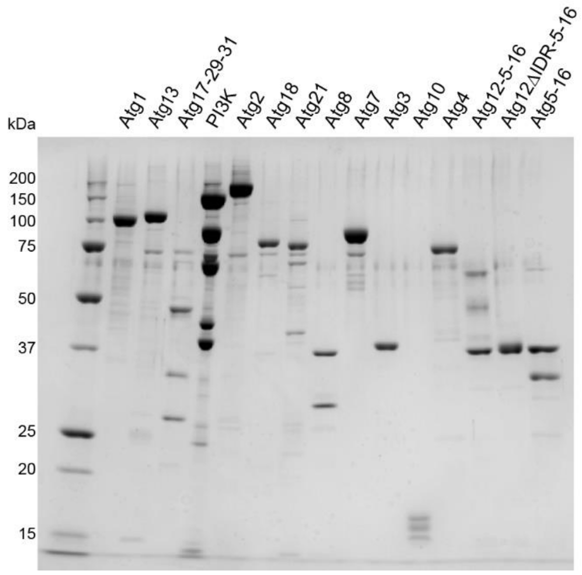


**Figure S1. SDS-PAGE image of the proteins used for Figure 1 experiments (related to Figure 1).**

Figure S2

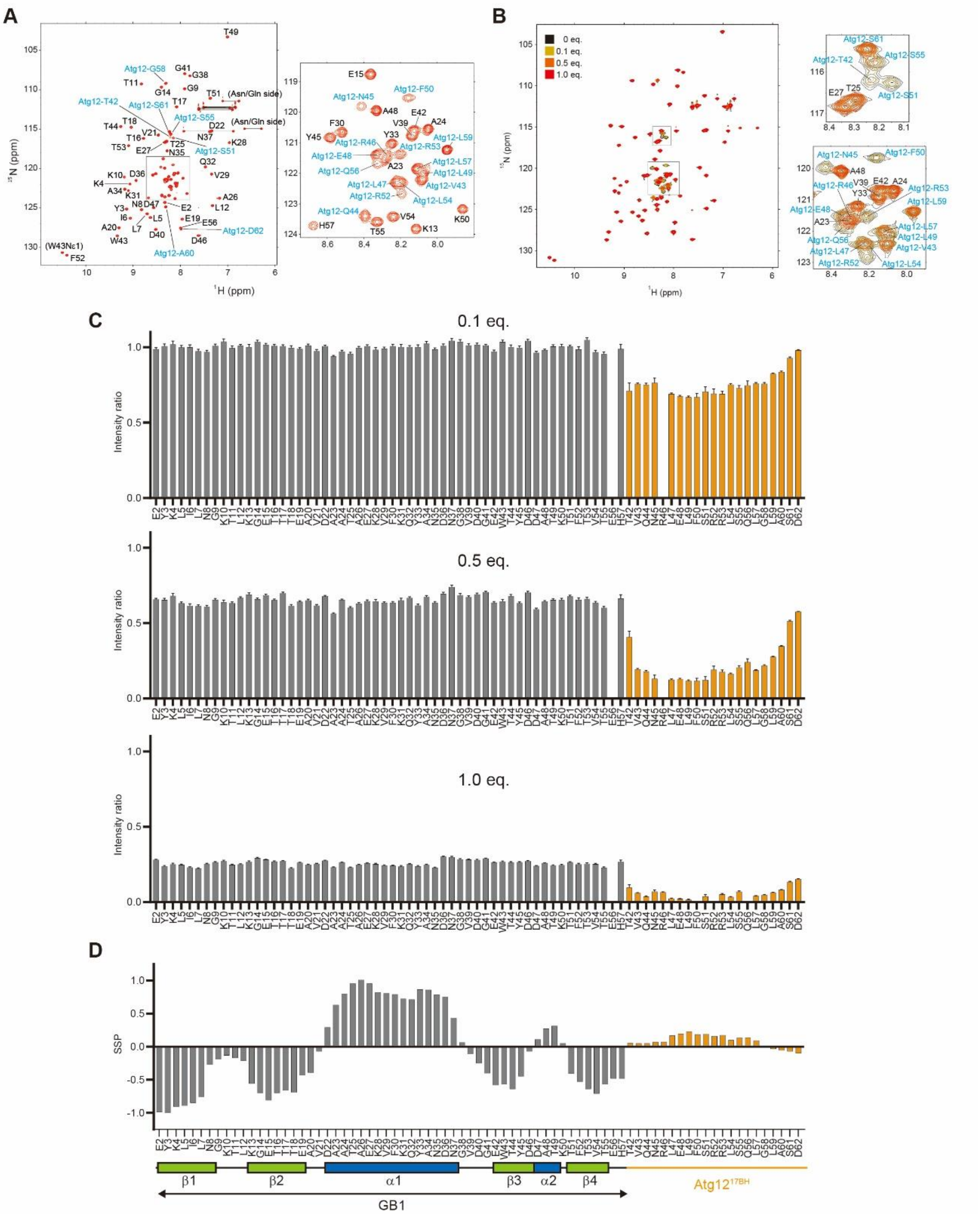


**Figure S2. NMR analysis of GB1–Atg12$^{17BH}$ (related to Figure 2).** (A) Main chain assignment results are shown on [$^{1}$H-$^{15}$N] heteronuclear single quantum coherence spectroscopy (HSQC); GB1-derived signals are labeled in black, and Atg12-derived signals are labeled in blue. (B) Superposition of [$^{1}$H-$^{15}$N] HSQC obtained by titration experiments. (C) Signal intensity ratios before and after the addition of Atg17–Atg31C are plotted by residue (GB1 is shown in gray and Atg12 in orange, 0.1, 0.5, and 1.0 molar equivalents from top to bottom). The signal-to-noise ratio of the signals was used to calculate the error bars. (D) Secondary structure propensity analysis results (GB1 is shown in gray and Atg12 in orange). The secondary structure position in the GB1 structure is attached.

**Figure S3**

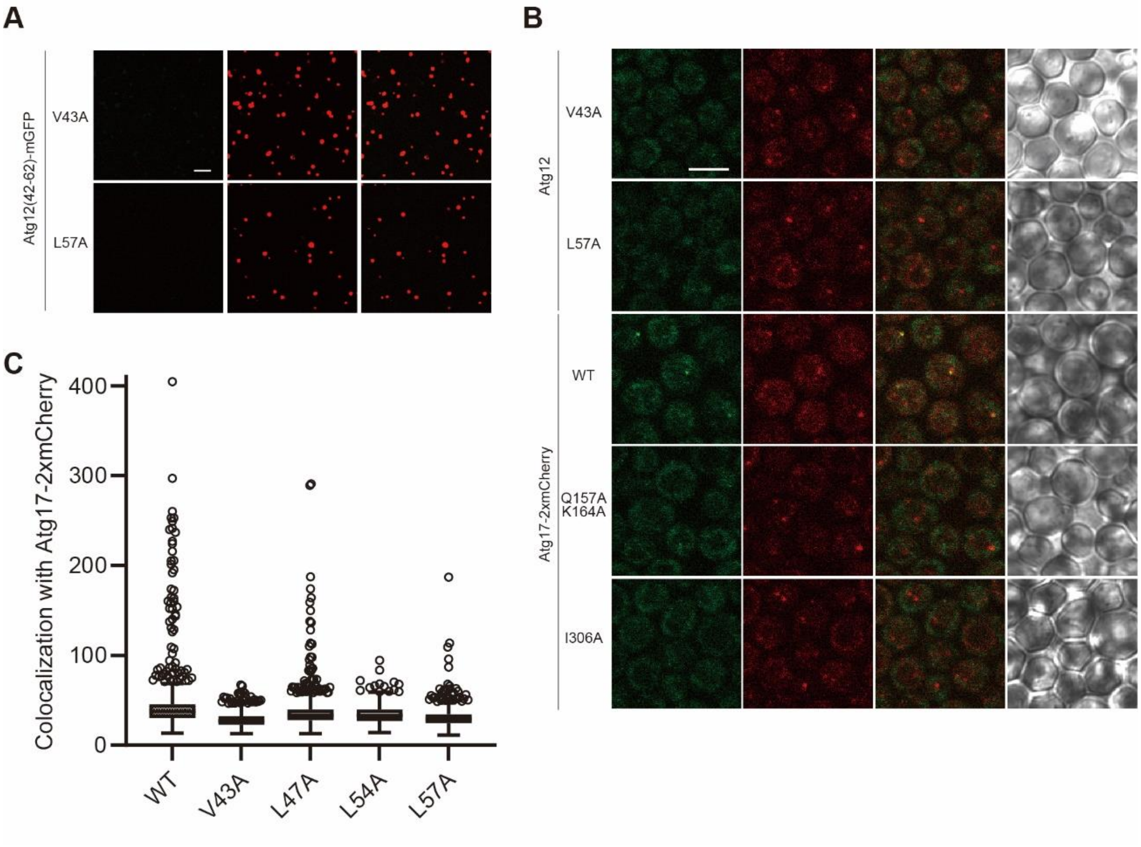


**Figure S3. Specific Atg12$^{17BH}$–Atg17 interaction is important for the condensation at the droplet both in vitro and in vivo (related to Figure 3).** (A) Condensation of GFP–Atg12$^{17BR}$ at the early PAS droplets was impaired by mutations. (B) PAS targeting of the Atg12–Atg5–Atg16 complex was impaired by mutations. (C) Overall view of Figure 3E.

Figure S4

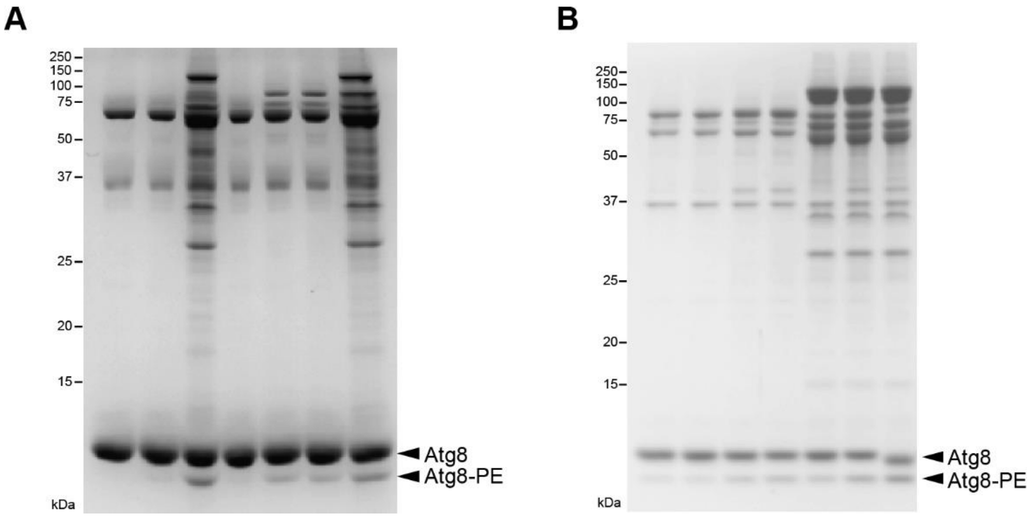


**Figure S4. Effect of the PAS droplets, Atg21, and PI3P on Atg8 lipidation (A) and delipidation (B) reactions (related to Figure 4).**

### Figure S5

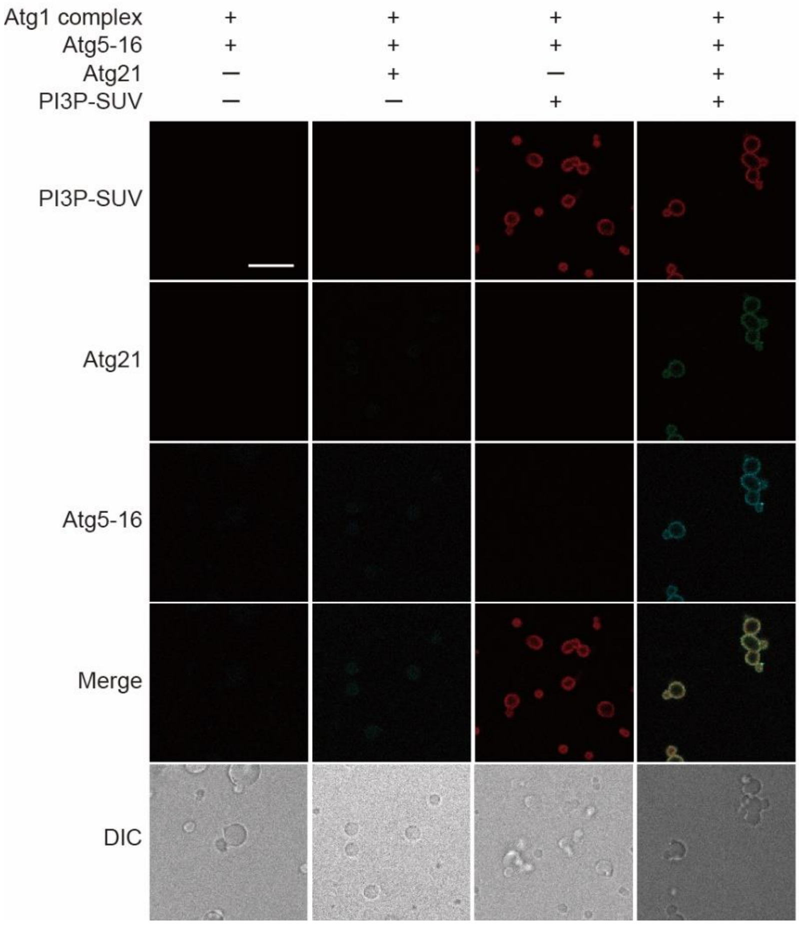


**Figure S5. In vitro Atg21 and PI3P-dependent targeting of the Atg5–Atg16 complex to the PAS droplets observed by fluorescence microscopy (related to Figure 5).** Scale bar = 10 μm.

#### Figure S6

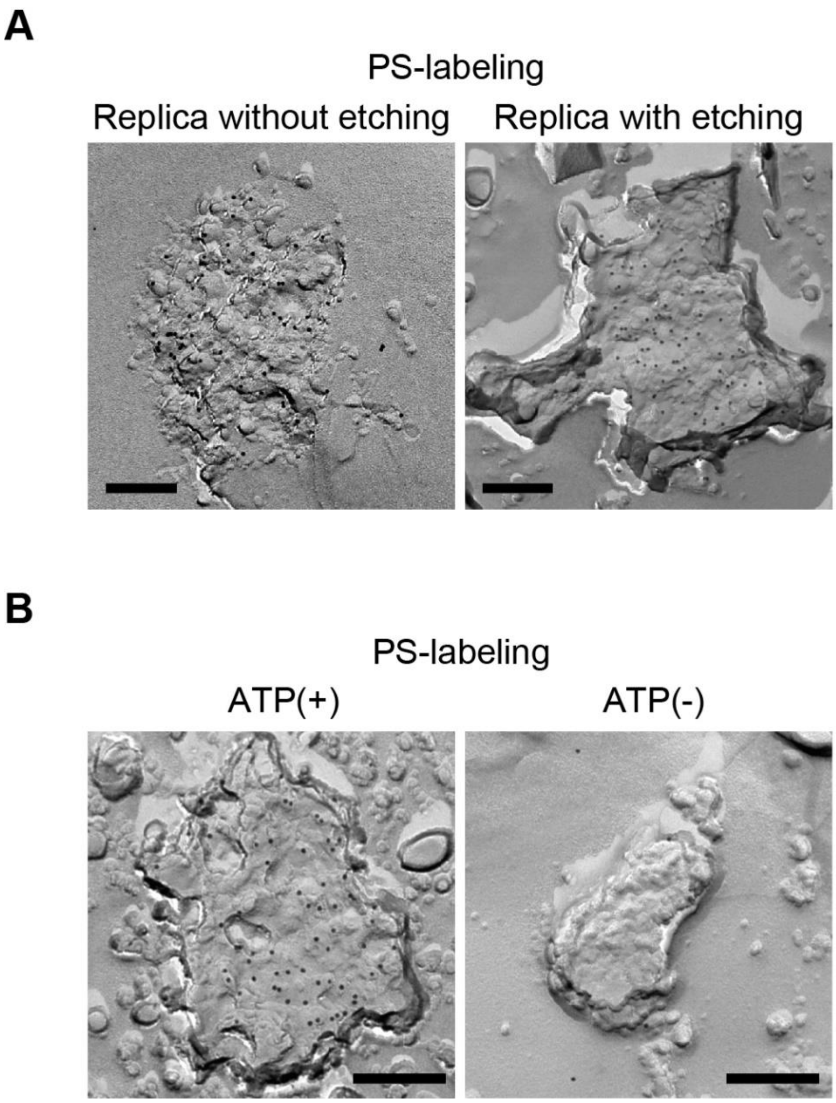


**Figure S6. Replica electron microscopy of the early PAS droplets (related to Figure 6).** (A) Replica electron microscopy with or without etching. (B) Replica electron microscopy in the presence of Atg21 but in the absence of Atg12–Atg17 interaction. Scale bar = 200 nm.

Table S1. Yeast strains used in this study.

| Strain | Genotype | Source |
|---|---|---|
| BJ2168 | *MATa leu2 trp1 ura3-52 prb1-1122 pep4-3 prc1-407 gal2* | Y.G.S.C. |
| W303-1A | *MATa ade2-1 ura3-1 his3-11,15 trp1-1 leu2-3,112 can1-100* | (Thomas & Rothstein, 1989) |
| ScTK1793 | BJ2168, *atg17*Δ::*natNT2* | This study |
| ScTK1806 | BJ2168, *ATG5–3×FLAG-kanMX4 atg17*Δ::*natNT2* | This study |
| ScTK1852 | W303-1A, *ade2-1*::*ADE2 ATG5-EGFP-kanMX4 atg17*Δ::*hphNT1 atg21*Δ::*zeoNT3 ATG17pro*::pRS305-*ATG17-2 × mCherry atg12*Δ::*natNT2 his3-11*::*pRS303-ATG12* | This study |
| ScTK1854 | W303-1A, *ade2-1*::*ADE2 ATG5-EGFP-kanMX4 atg17*Δ::*hphNT1 atg21*Δ::*zeoNT3 ATG17pro*::pRS305-*ATG17-2×mCherry atg12*Δ::*natNT2 his3-11*::*pRS303-atg12*$^{L47A}$ | This study |
| ScTK1855 | W303-1A, *ade2-1*::*ADE2 ATG5-EGFP-kanMX4 atg17*Δ::*hphNT1 atg21*Δ::*zeoNT3 ATG17pro*::pRS305-*ATG17-2×mCherry atg12*Δ::*natNT2 his3-11*::*pRS303-atg12*$^{L54A}$ | This study |
| ScTK566 | W303-1A, *ade2-1::ADE2 PGK1-EGFP-kanMX4 atg12*Δ::*natNT2* | This study |
| ScTK570 | W303-1A, *ade2-1*::*ADE2 PGK1-EGFP-kanMX4 atg12*Δ::*natNT2 atg21*Δ::*zeoNT3* | This study |

Y.G.S.C. (Yeast Genetic Stock Center)
